# Investigating the impact of introduced crabs on the distribution and morphology of littorinid snails: Implications for the survival of the snail *Littorina saxatilis*

**DOI:** 10.1101/2023.01.28.526005

**Authors:** Christopher D. Wells, Kaitlin S. Van Volkom, Sara Edquist, Sinead Marovelli, John Marovelli

**Author notes:** Christopher Wells and Kaitlin Van Volkom contributed equally to this study. First author was decided by a coin flip.

## Abstract

Introduced species can have profound impacts on communities by displacing and consuming native species. The intertidal communities in the Gulf of Maine have been radically altered through a suite of invasions including the herbivorous snail *Littorina littorea* and the omnivorous crab *Carcinus maenas* leading to morphological and spatial distribution changes in two native gastropod grazers (*Littorina saxatilis* and *Littorina obtusata*). Subsequently, another intertidal omnivorous crab *Hemigrapsus sanguineus* has become abundant in the intertidal, occurring in areas which were once refuges from crab predation. In order to quantify the potential impact of *H. sanguineus* on native snails, we conducted both *in* and *ex situ* experiments, testing the susceptibility of different sized snails to predation by introduced crabs. Additionally, we measured the distribution, abundance, and morphology of intertidal snails and crabs. Smaller snails of all species were the most susceptible to predation, although susceptibility differed among snail species. *Littorina saxatilis* was the most vulnerable to predation, with 73 and 64% of the population susceptible to large *H. sanguineus* and small *C. maenas*, respectively, while more than 89% of the *L. littorea* population was resistant to predation. *Littorina saxatilis* has been relegated to the high intertidal where there is high abiotic stress and poor-quality food, but until the invasion of *H. sanguineus*, there was little predation risk. Now, it seems that *L. saxatilis* is at the most risk of all three snail species, and may be at threat of local extirpation at locations with high populations of *H. sanguineus.*

## 1. Introduction

The Gulf of Maine is a low diversity system (Witman et al., 2004) historically influenced by top-down control from large predatory fishes (Jackson et al., 2001; Steneck et al., 2004). Declines in large predators, rapid warming, and shifting seasonality have facilitated sweeping changes in local communities (Pershing et al., 2015; Staudinger et al., 2019). In addition, the Gulf of Maine has a long history of species introductions, which alter community structure through competition and predation with native species (Carlton, 2003). Many benthic communities are now dominated by introduced algae and invertebrates, which have changed the demersal invertebrate community (Dijkstra et al., 2017; Valentine et al., 2007). In the intertidal, the algal assemblage is partly dictated by their palatability to an introduced littorinid snail (Lubchenco, 1978).

Prior to European colonization, the rocky intertidal in the Gulf of Maine was dominated by two species of native littorinid snails, *Littorina saxatilis* Olivi, 1792 *and Littorina obtusata* Linnaeus, 1758 (Blakeslee et al., 2008; Panova et al., 2011). Currently, the dominant littorinid snail is the introduced periwinkle *Littorina littorea* Linneaus, 1758, which has been present in New England since at least the 19^th^ century, likely coming over in ballast (Carlton, 1982). Littorina *littorea* is a generalist grazer, preferring periphyton, germlings, and algae without chemical or structural defenses (reviewed in Reid, 1996). The invasion of *L. littorea* and the subsequent competition for food has relegated *L. saxatilis* to the high intertidal and supralittoral zone (Behrens Yamada and Mansour, 1987; Reid, 1996), where food resources are low and abiotic stressors such as desiccation, temperature variability, and freezing risk is high. The other native littorinid, *L. obtusata*, seems to be unaffected by *L. littorea*, likely because *L. obtusata* is a fucoid specialist (Watson and Norton, 1987), which *L. littorea* only consumes as germlings.

One of the main predators of littorinid snails in New England is introduced crabs (reviewed in Teck, 2006). Prior to European colonization, the prominent large crustacean predators in New England were two cancrid crabs *Cancer borealis* Stimpson, 1859 and *Cancer irroratus* Say, 1817 and the American lobster *Homarus americanus* H. Milne Edwards, 1837. These three species forage throughout the intertidal, but are restricted by the tide; highest levels of predation occur within the low intertidal (Ellis et al., 2007). While these species are capable of consuming littorinid snails (Donahue et al., 2009; Jones and Shulman, 2008), snails were rarely found in field-collected gut contents (e.g., Donahue et al., 2009; Stehlik, 1993). The introduced European green crab *Carcinus maenas* Linnaeus, 1758, which first appeared in the Gulf of Maine in the 19^th^ century, is an aggressive, generalist predator, consuming mollusks, crustaceans, polychaetes, and algae (Ropes, 1968), and quickly became the dominant crustacean in the middle and lower intertidal (reviewed in Teck, 2006). More recently, the Asian shore crab *Hemigrapsus sanguineus* De Haan, 1835 was introduced in 1988, and reached the Gulf of Maine in 2000 (Bourdeau and O’Connor, 2003), spreading as far North as Schoodic Peninsula, Maine (Delaney et al., 2008). Since its introduction, the abundance of *H. sanguineus* has rapidly increased (Kraemer et al., 2007), and in some regions has displaced *C. maenas* as the dominant intertidal crustacean predator (Griffen et al., 2008). *H. sanguineus* can survive at all tidal heights including the high intertidal zone which once served as a gastropod refuge from *C. maenas* (Eastwood et al., 2006). This shift in predation pressure could have large impacts on littorinid populations, which may alter gastropod grazing pressure on local algal communities.

In this study, we investigate the relationships between both introduced crab species and the three littorinid snails in the Gulf of Maine. Densities and morphological measurements of crabs and snails at three locations across tidal heights and season on the coast of New Hampshire and Southern Maine are reported. Using snail shell morphology measurements, we determine critical changes in shell investment as snails grow. Additionally, we examine the impacts of predation on littorinid populations with both *in situ* and *ex situ* experiments to determine snail predation susceptibility to each crab. This study represents a large effort to quantify and describe crab and gastropod presence in the intertidal and investigate the trophic relationships between these species.

## 2. Materials and Methods

### 2.1. Study Sites

Three locations along the coast of the Gulf of Maine were sampled for littorinid and crab densities and morphological parameters: Seapoint Beach, Kittery Point, Maine (43.0909 °N, 70.6600 °W); Fort Stark, New Castle, New Hampshire (43.0584 °N, 70.7114 °W); and Odiorne Point, Rye, New Hampshire (43.0454 °N, 70.7130 °W). All sites were exposed rocky intertidal communities dominated by boulder and ledge habitat with neighboring areas of sandy beach and protected coves. All sites exhibited distinct zonation typical of the Gulf of Maine (Lubchenco, 1980). In particular, the high intertidal zone was characterized by large cyanobacteria-covered boulders, the mid-intertidal zone was characterized by the brown fucoid algae *Ascophyllum nodosum* (Linnaeus) De Jolis, 1863 and *Fucus* spp., the low intertidal zone was characterized by red algae, and the shallow subtidal was characterized by coralline algae and kelps, in addition to introduced algae (Dijkstra et al., 2017).

### 2.2. Field Surveys

To determine the vertical distribution, size structure, and abundance of the three littorinid species (*Littorina littorea*, *Littorina obtusata*, and *Littorina saxatilis)* and the four most common decapod crab predators (*Carcinus maenas*, *Hemigrapsus sanguineus*, *Cancer borealis*, and *Cancer irroratus*), a series of seasonal surveys were conducted at each site. Sampling was conducted in December 2011; March, June, September, and December 2012; and June 2016. During December 2011, only Seapoint Beach and Odiorne Point were sampled. Surveys were conducted within the shallow subtidal and three intertidal heights, defined by their algal composition (see above). Tide pools were excluded from sampling, as they represent distinct communities (Lubchenco, 1978).

Quadrats were placed randomly along a 100 m transect line at each tidal height. Quadrat location along the transect line was determine by random number generator; if random numbers repeated, the area was resampled. To determine the density of littorinid populations, all snails were counted within ten 0.25 m^2^ quadrats at each height. All *Littorina spp.* within a 0.0625 m^2^ quadrat, nested within the larger 0.25 m^2^ quadrat, were collected for morphological characterization. When logistically possible we collected a minimum of ten individuals per species for each intertidal zone. Due to logistical limitations, only ten quadrats were collected in the shallow subtidal. Shell height, aperture length, and shell thickness were measured with digital calipers for each snail. To determine the density and size structure of crabs, all *C. maenas*, *H. sanguineus*, *C. irroratus*, and *C. borealis* were collected within ten 1.0 m^2^ quadrats at each height. All crabs were transported to the lab where width and length of the larger cheliped and carapace width were measured and sex was determined. Crabs that were not presenting external morphological characters characteristic of sexual mature crabs (i.e., modified abdomen) were identified as juvenile crabs.

### 2.3. Field Feeding Trials

To determine the feeding preferences of *H. sanguineus* and *C. maenas*, we conducted *in-situ* experiments in August and September of 2012 at Fort Stark. Snails were enclosed in cages with one of three types of crab or no crab (control). The crab treatments were either one 35-mm carapace width *H. sanguineus,* one 20-mm carapace width *H. sanguineus*, or one 35-mm carapace width *C. maenas*. All crabs were male and had two fully-grown and intact claws. Crab sizes were informed by the initial surveys. 35-mm carapace width *H. sanguineus* are relatively large individuals within our survey area, 20-mm carapace width *H. sanguineus* are average sized individuals, and 35-mm carapace width *C. maenas* were used for comparison with the largest *H. sanguineus*. Eleven snails were placed within each cage, divided into 5 mm shell height size class ranges. We used three size classes for *L. obtusata* and *L. saxatilis* (2.5-5 mm, 5.01-10 mm, and 10.01-15 mm) and five size classes for *L. littorea* (2.5-5 mm, 5.01-10 mm, 10.01-15 mm, 15.01-20 mm, and 20.01-25 mm). Maximum size classes were determined by availability of snails.

All animals were haphazardly collected from Fort Stark within one week of deploying the cages and maintained in the laboratory at 10 °C in 32 ppt seawater until deployment. All snails had intact shells with no large chipping or scars. Cages were constructed from 1-cm square-opening polyvinyl chloride coated steel mesh with a 1 mm plastic mesh cage nested within. Each cage was 15 × 15 × 5 cm. Sets of four cages (one from each treatment) were attached together with hog rings to reduce the effect of spatial heterogeneity on crab behavior. At each of three intertidal heights two sets of cages were secured and buried under small boulders to simulate natural habitat and shelter. After 14 days, all cages were retrieved, and presence or absence of each snail was documented. All damage was scored by the same investigator to ensure consistent scoring across all trials. Three replicate deployments were conducted for a total of 18 replicates per treatment. However, during the third deployment, Hurricane Sandy swept away the two middle intertidal blocks. An additional four cages within sets of four were torn open, allowing for snails to escape the cages, and were therefore not used in analyses.

### 2.4 Laboratory Feeding Trials

To determine the potential susceptibility to the smallest sized snails, we conducted a laboratory experiment using both *C. maenas* and *H. sanguineus* and all three snail species in the summer of 2016. Snails and crabs were collected haphazardly from Fort Stark and held at 10 °C for three to five days. During this time, food was withheld from crabs. Individual crabs were placed in plastic 19.4 ×16.5 ×11.4 cm containers and maintained at 10 °C. During each trial, crabs were given 18 snails of one species, evenly distributed among nine size categories: 2.0-2.9. 3.0-3.9, 4.0-4.9, 5.0-5.9, 6.0-6.9, 7.0-7.9, 8.0-8.9, 9.0-9.9, and 10.0-12.0. Crabs were given 24 hours to feed and then the number of snails consumed was recorded. Each crab was fed all three snail species during the course of the experiment in three separate trials. In between trials, crabs were fed mussels and then starved for three days to standardize hunger levels. The order of the snail species fed to each crab was randomized.

### 2.5 Statistical analysis

All statistical analyses were performed in R version 4.2.2 (R Core Team, 2022).

Relationships between snail shell morphology parameters were assessed with Davies’ tests and change-point regression models (R package segmented 1.6-2, Muggeo, 2003). Davies’ test can test for a non-constant relationship between two shell parameters. Change-point regression models allow for quantitative estimation of segmented relationships between two morphology parameters and indicate the change-point value (i.e., the point of inflection). When a segmented linear relationship was not found, a linear regression was performed to relate the two parameters. Segmented relationships were allowed to have one breakpoint and 50 bootstrap samples were used in the bootstrap restarting algorithm.

Relationships between snail shell morphology parameters (i.e., shell length, aperture length, and shell thickness) and tidal height, sample season, and their interaction were evaluated with linear mixed effects models (LMM, R package nlme 3.1-160, Pinheiro et al., 2022). Linear mixed effect models for crab morphology parameters (i.e., carapace width, cheliped length, and cheliped width) also had sex as an additional fixed factor and no interactive effects were modeled. For both groups, site and quadrat within site were modeled as random factors. Tidal heights which had less than 20 individuals of a species were excluded from the analyses and if a species only occurred at one tidal height, height was dropped from the model.

The impact of season, height, and their interactive effect on snail and crab densities were evaluated with generalized linear mixed effect models with Poisson error distributions (GLMM, R package lme4 1.1-31, Bates et al., 2015). Models were tested for over- and underdispersion and where models were over- or underdispersed, negative binomial error distributions were used. Site was incorporated into the model as a random factor. Where models were nearly unidentifiable (unstable fitting of model parameters) interactive effects were not evaluated. Negative binomial linear mixed effect models perform poorly with inflated-zero data and so when a tidal height had a mean of less than 1 individual × m^−2^, that tidal height was dropped.

For both the laboratory and field predation experiment, the effect of crab treatment, snail species, and size of snail on survival of snails were evaluated with generalized linear models (GLM) with binomial error distributions. All interactive effects were evaluated. Population-level risk of predation by crab treatments was assessed by taking the sum of the products of the proportion of snails at a given shell length and the predicted proportion that would survive in an interaction with a crab derived from the binomial GLM.

## 3. Results

### 3.1 Density and morphometric surveys

Across sampling seasons, sites, and tidal heights, 5943 *Littorina littorea*, 1683 *Littorina obtusata*, and 1760 *Littorina saxatilis* were measured for shell morphology parameters. An additional 10,260 *L. littorea*, 3410 *L. obtusata*, and 1021 *L. saxatilis* were counted for density estimates. For crabs, 2189 *Hemigrapsus sanguineus*, 1132 *Carcinus maenas*, 130 *Cancer irroratus*, and 48 *Cancer borealis* were measured for carapace and cheliped morphology parameters and density estimates.

Crabs and snails had clear preferences for specific tidal heights and generally those preferences were dependent on the season (Fig. 1). Density of all snail species and *H. sanguineus* had a significant interactive response to sampling season and tidal height (GLMM, X^2^ ≥ 19.1, df = 3-9, *p* < 0.01), but each pattern was species specific. *C. maenas* had insufficient densities to evaluate the interactive effect of sampling season and tidal height but both factors separately had significant effects on *C. maenas* distribution (GLMM, X^2^ ≥ 60.3, df = 2-3 *p* < 0.01). Both *Cancer* spp. had insufficient data for analyzing the effects of tidal height and season on density but densities across these factors are displayed in Figure 1.

**Fig. 1.**
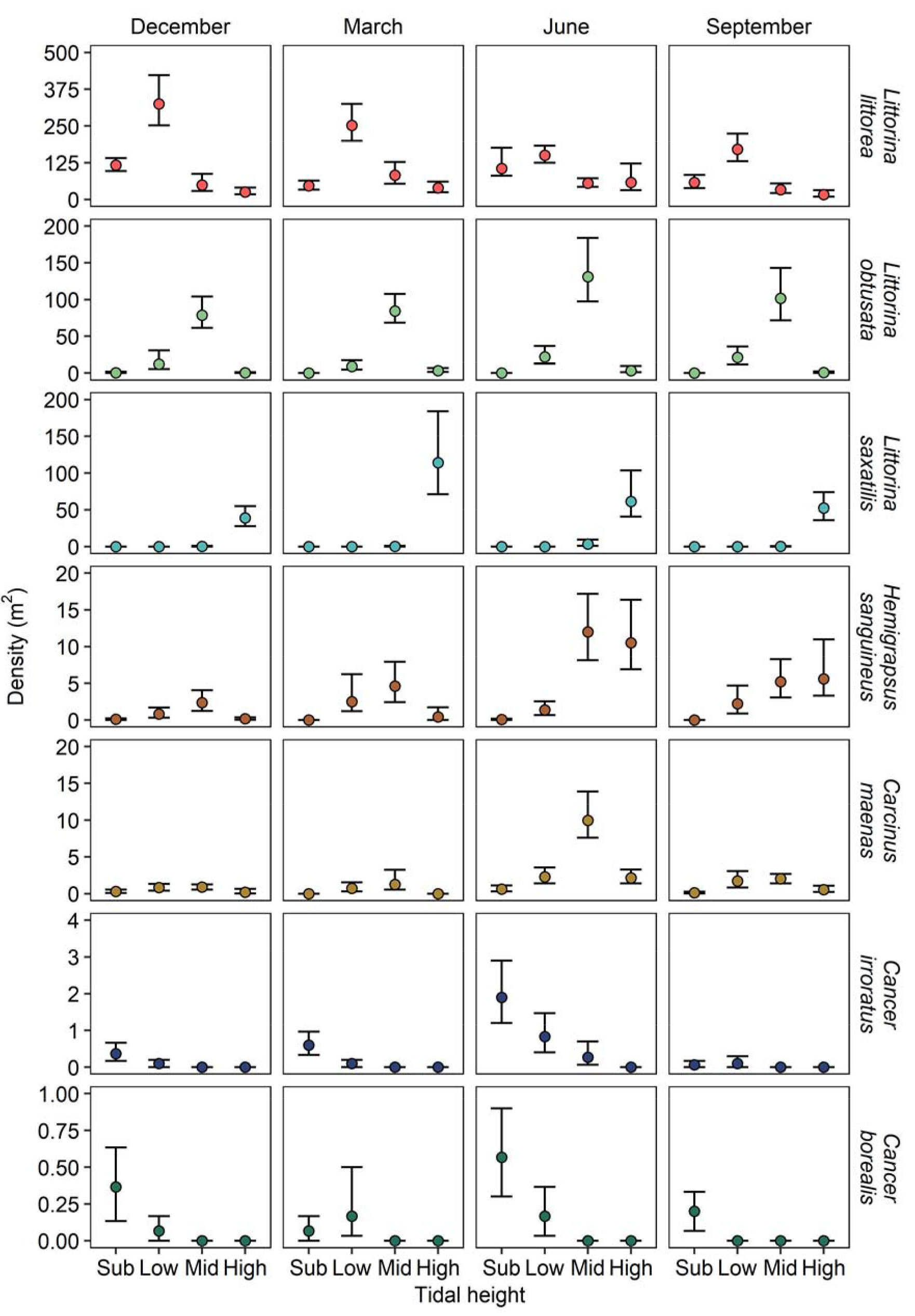
Mean density of intertidal and shallow subtidal littorinid snails and crabs across tidal heights and sampling seasons. Error bars are 95% bootstrapped confidence intervals.

*L. littorea* was present in every tidal height and during every sampling season and was the most abundant snail at the sites we sampled. *Littorina littorea* was most abundant in the lower intertidal (mean of 216 snails × m^−2^) and during June and December (85.0 and 81.1 snails × m^−2^, respectively) and least abundant in the upper intertidal (30.5 snails × m^−2^) and during September (48.6 snails × m^−2^). *Littorina littorea* was most evenly distributed across tidal heights during the warmer months (June and September) and had a strong preference for the lower intertidal during the colder months (December and March). *Littorina littorea* shell length, aperture length, and shell thickness had significant interactive responses to sampling season and tidal height (Fig 2, LMM, *F*_9, 618_ ≥ 4.1, *p* < 0.01). *Littorina littorea* were largest during March (mean of 19.6 mm shell length) and in the subtidal (24.2 mm) and smallest during December (17.8 mm) and the upper intertidal (12.8 mm).

**Fig. 2.**
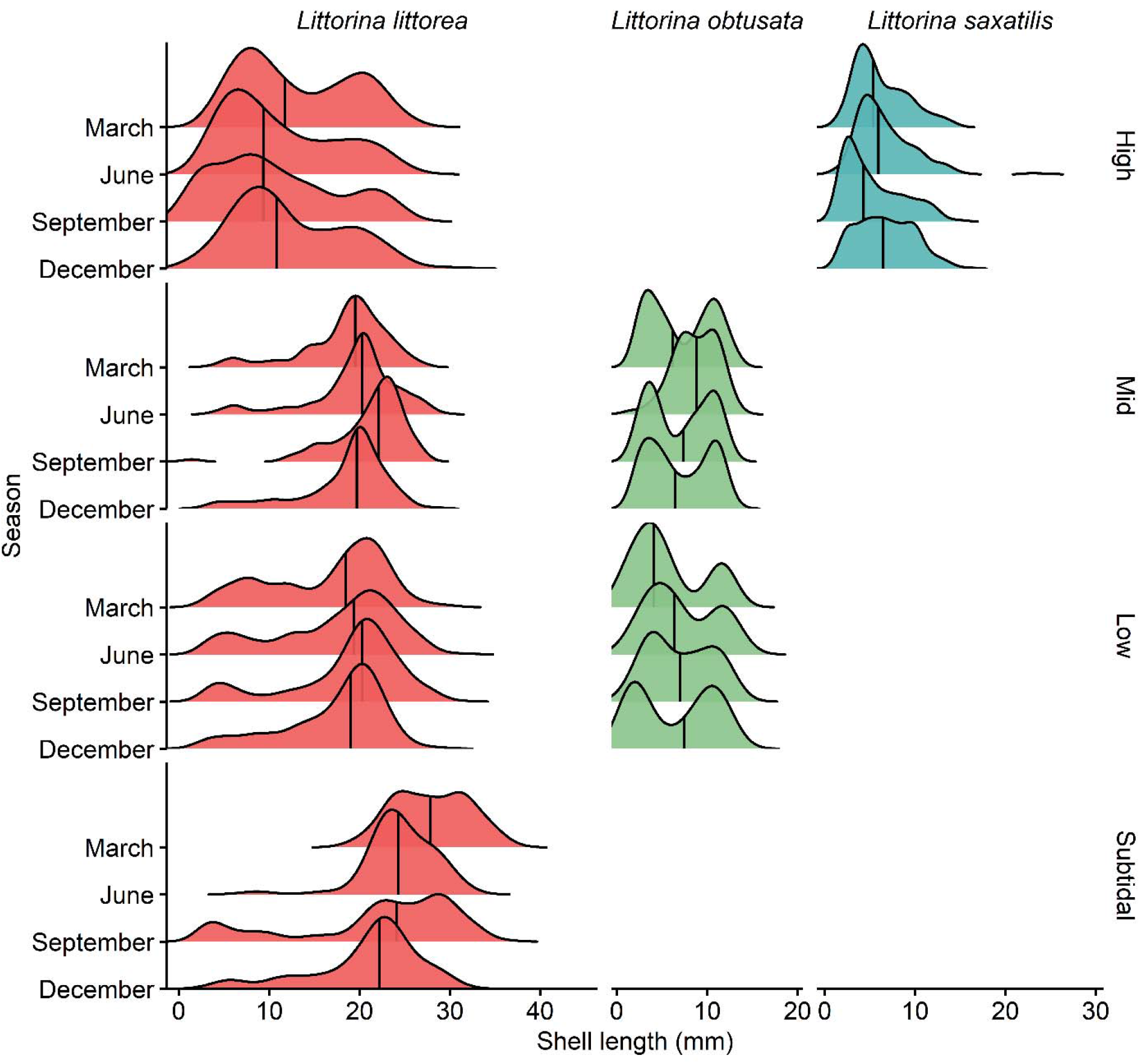
Size-frequency distribution of littorinid snails across tidal heights and seasons. Solid vertical lines indicate median shell length.

*Littorina obtusata* was primarily found in the lower and middle intertidal zones (Fig. 2), almost exclusively associated with fucoid algae (e.g., *Ascophyllum nodosum* and *Fucus distichus* Linnaeus, 1767). Several individuals were found in the subtidal (6 individuals) and upper intertidal (85) and were either associated with particularly high elevation *Fucus* spp. or seemed to have been tossed into the location by wave action. *Littorina obtusata* was most abundant in the middle intertidal (mean of 97.5 snails × m^−2^) and during June (18.8 snails × m^−2^) and least abundant in the subtidal (0.2 snails × m^−2^) and during December (6.1 snails × m^−2^). *Littorina obtusata* aperture length had a significant interactive response to sampling season and tidal height (Fig 2, LMM, *F*_3, 257_ = 3.1, *p* = 0.03). Shell length did not have a significant interactive response to sampling season and tidal height (Fig 2, LMM, *F*_3, 257_ = 2.4, *p* = 0.07) nor the main effect of tidal height (Fig 2, LMM, *F*_1, 257_ = 2.5, *p* = 0.11), but was significantly related to sampling season (Fig 2, LMM, *F*_3, 257_ = 6.7, *p* < 0.01). Shell thickness also did not have a significant interactive response to sampling season and tidal height (Fig 2, LMM, *F*_3, 257_ = 1.6, *p* = 0.19), but did significantly respond to the main effect of tidal height (Fig 2, LMM, *F*_1, 257_ = 15.7, *p* < 0.01) and did not significantly respond to sampling season (Fig 2, LMM, *F*_3, 257_ = 0.8, *p* = 0.51). *Littorina obtusata* was smallest in March (mean of 7.0 mm) and largest in June (8.6 mm).

*Littorina saxatilis* were almost exclusively found in the upper intertidal (mean of 60.6 snails × m^−2^), taking refuge in moist gravel, under rock overhangs, and within rock crevices (Fig. 2). Across all locations and seasons, only 61 individuals were found in the middle intertidal and none were found in the lower intertidal or subtidal. *Littorina saxatilis* was most abundant in June (mean of 12.8 snails × m^−2^) and least abundant in December (3.0 snails × m^−2^). Sufficient *L. saxatilis* were collected only in the high intertidal for shell morphology analyses and therefore the effect of tidal height on shell parameters was not evaluated. Sampling season did not significantly affect shell and aperture length of *L. saxatilis* (Fig 2, LMM, *F*_3, 187_ ≤ 2.5, *p* ≥ 0.06), but did significantly affect shell thickness (Fig 2, LMM, *F*_3, 187_ = 4.8, *p* < 0.01).

*Hemigrapsus sanguineus* was the most abundant crab we found during our surveys and occurred at all sampled tidal depths (Fig. 3B), although few were found in the subtidal (5 individuals). They were most abundant in the middle intertidal (mean of 3.3 crabs × m^−2^) and during June (3.8 crabs × m^−2^) and least abundant in the subtidal (0.04 crabs × m^−2^) and during December (0.6 crabs × m^−2^). Carapace width and cheliped length and width were significantly related to season (Fig 3A, LMM, *F*_3, 135_ ≥ 8.9, *p* < 0.01), height (Fig 3B, LMM, *F*_2, 135_ ≥ 3.8, *p* < 0.01), and sex (Fig 3C, LMM, *F*_2, 1031_ ≥ 124, *p* < 0.01). Crabs were largest in December (mean of 16.0 mm carapace width), in the lower and upper intertidal (15.1 mm), and males were the larger sex (18.4 mm).

**Fig. 3.**
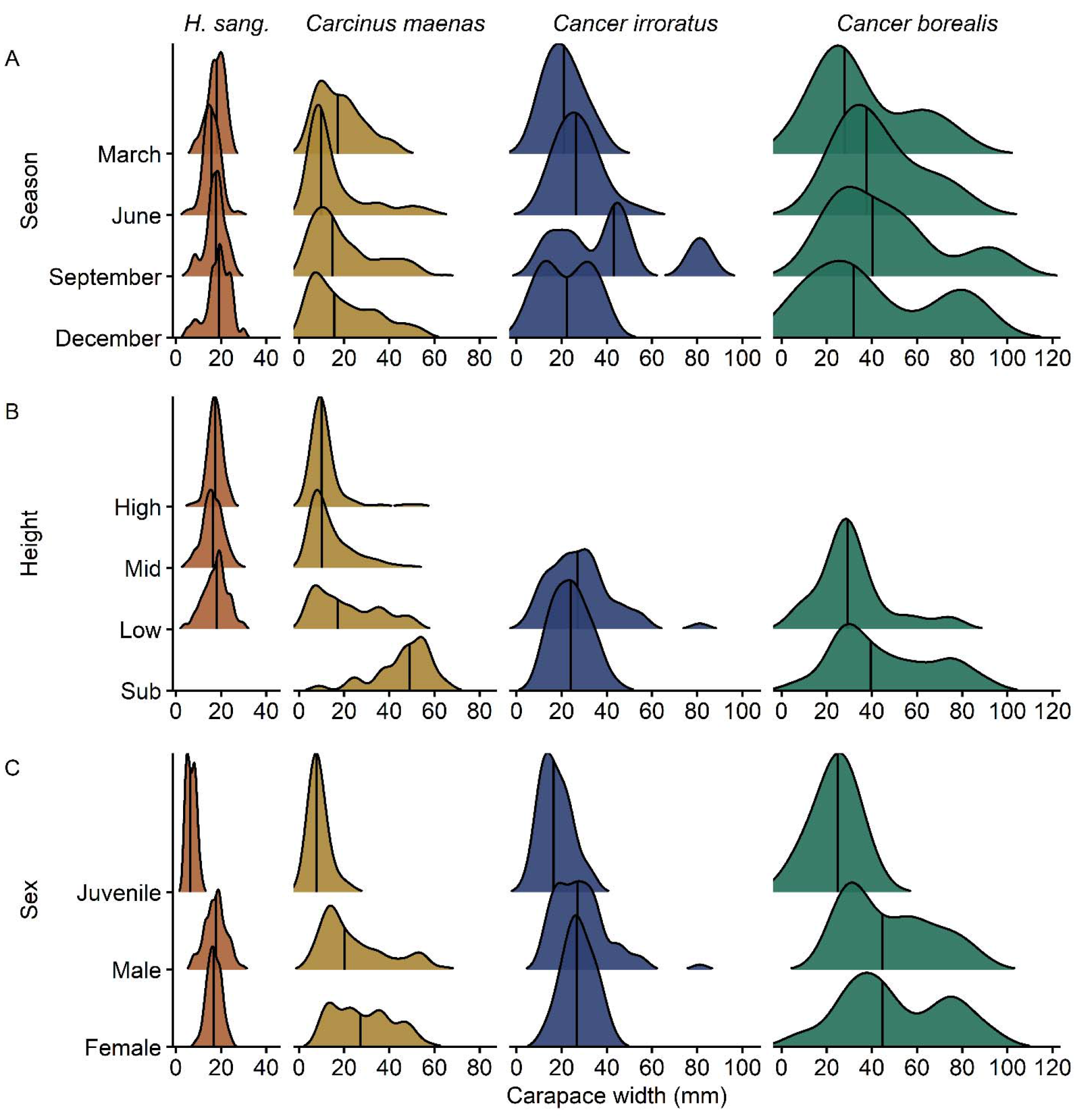
Size-frequency distribution of crabs across (A) seasons, (B) tidal heights and (C) sex. Solid vertical lines indicate median carapace width. Distributions are not displayed for tidal heights and seasons that had fewer than 10 crabs.

*C. maenas* also occurred at all sampled tidal depths. They were most abundant in the middle intertidal (mean of 2.3 crabs × m^−2^) and during June (3.9 crabs × m^−2^) and least abundant in the subtidal (0.3 crabs × m^−2^) and during March (0.5 crabs × m^−2^). Carapace width and cheliped length and width were significantly related to season (Fig 3A, LMM, *F*_3, 177_ ≥ 2.7, *p* < 0.05), height (Fig 3B, LMM, *F*_2, 177_ ≥ 98.9, *p* < 0.01), and sex (Fig 3C, LMM, *F*_2, 351_ ≥ 138, *p* < 0.01). Crabs were largest in March (mean of 28.9 mm carapace width), in the subtidal (42.1 mm), and females were the larger sex (32.0 mm).

*C. borealis* was rare, occurring only in the subtidal and lower intertidal. They were most abundant in the subtidal (mean of 0.3 crabs × m^−2^) and during June (0.2 crabs × m^−2^). Carapace width and cheliped measurements were significantly related to height (Fig 3A, LMM, *F*_1, 33_ ≥ 4.7, *p* ≤ 0.04) and sex (Fig 3C, LMM, *F*_2, 10_ ≥ 9.7, *p* < 0.01), but not season (Fig 3B, LMM, *F*_3, 33_ ≤ 0.5, *p* ≥ 0.68). Crabs were largest in the subtidal (mean of 46.6 mm carapace width) and females were the larger sex (50.2 mm).

*C. irroratus* was more abundant than *C. borealis* and had a broader distribution across the intertidal, occurring in the middle intertidal in low numbers (9 individuals). Like *C. borealis*, *C. irroratus* was most abundant in the subtidal (mean of 0.7 crabs × m^−2^) and during June (0.8 crabs × m^−2^). Carapace width and cheliped measurements were significantly related to season (Fig 3A, LMM, *F*_3, 62_ ≥ 6.2, *p* < 0.01) and sex (Fig 3C, LMM, *F*_2, 63_ ≥ 7.9, *p* < 0.01), but not height (Fig 3B, LMM, *F*_1, 62_ ≤ 2.9, *p* ≥ 0.09). Crabs were largest during September (mean of 38.4 mm carapace width) and females were the larger sex (31.8 mm).

Generally, regression models with segmented relationships fit better than linear models for nearly all species and combinations of shell metrics (Fig. 4, Table 1, Davies’ Test, *p* < 0.01). Only the relationship between *L. saxatilis* shell length and shell thickness fit better with a linear model (Fig. 2H, Table 1, Davies’ Test, *p* = 0.55). In both *L. littorea* and *L. obtusata*, investment in aperture lengthening halved at the change-point and thickening of the shell nearly tripled in *L. littorea* and more than tripled in *L. obtusata* (Table 1). While *L. saxatilis* had some significant change-points (i.e., between shell length and aperture length and between aperture length and shell thickness), they were less pronounced than either of the other littorinids (Table 1).

**Fig. 4.**
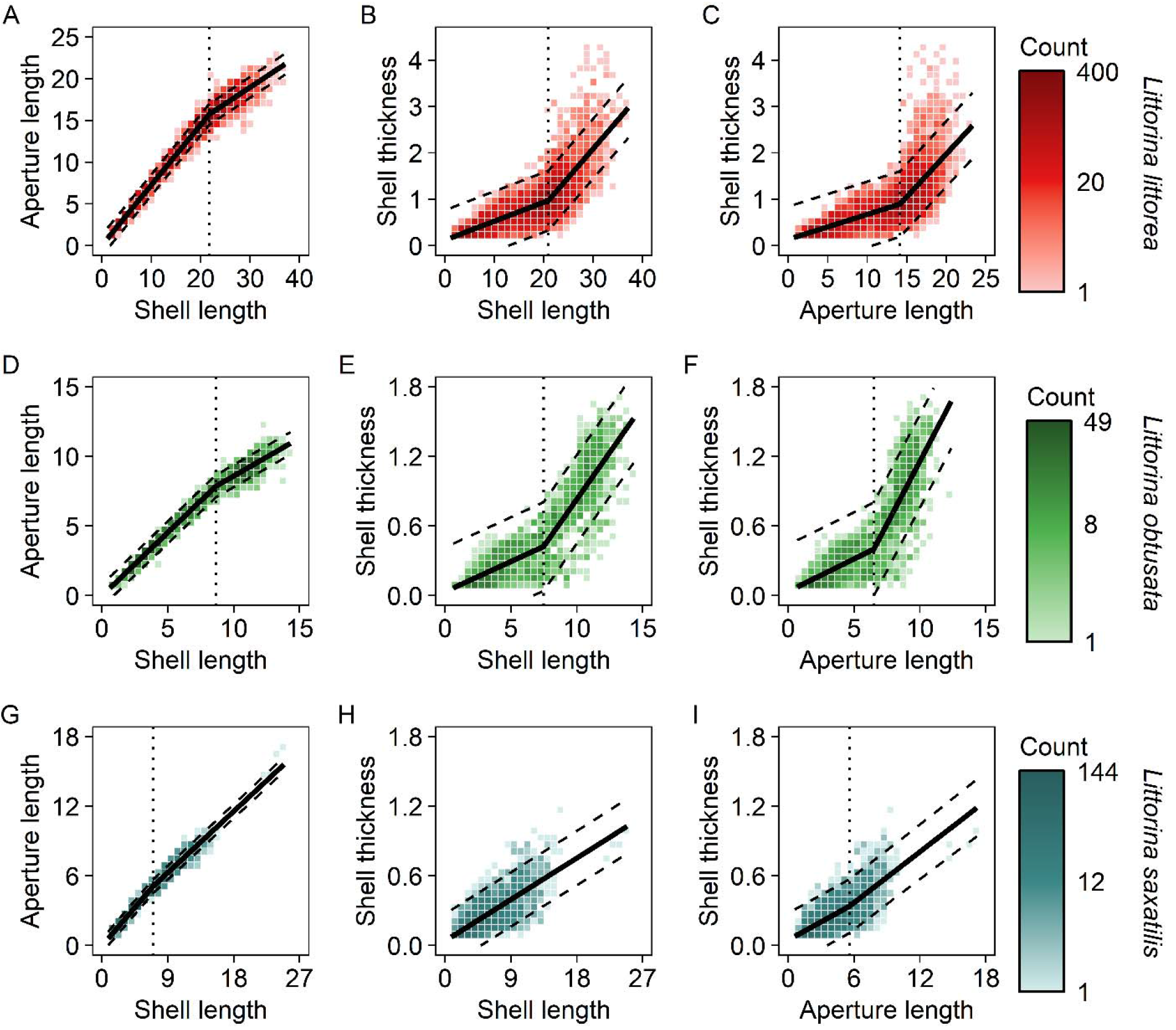
The relationship between (A, D) shell length and aperture length, (B, E) shell length and aperture thickness, and (C, F) aperture length and aperture thickness in the snails (A-C) *Littorina littorea*, (D-F) *Littorina obtusata*, and (G-I) *Littorina saxatilis*. Solid lines are change-point regression model fits, where significant change-points were detected, or linear regression fits with a 95% prediction interval (dashed lines). Dotted vertical lines indicate a significant change in slope.

**Table 1.** Coefficients and change-points for each regression model. A second coefficient and the change-point are reported where a regression model with segmented relationships fit better than a linear model.

| Species | Shell metric 1 | Shell metric 2 | Coefficient<br>1 | Coefficient<br>2 | Breakpoint<br>(mm) |
| --- | --- | --- | --- | --- | --- |
| *L. littorea* | Shell length | Aperture length | 0.72 | -0.33 | 21.7 |
| *L. littorea* | Shell length | Aperture thickness | 0.041 | 0.083 | 20.9 |
| *L. littorea* | Aperture length | Aperture thickness | 0.054 | 0.13 | 14.2 |
| *L. obtusata* | Shell length | Aperture length | 0.91 | -0.36 | 8.6 |
| *L. obtusata* | Shell length | Aperture thickness | 0.052 | 0.11 | 7.5 |
| *L. obtusata* | Aperture length | Aperture thickness | 0.056 | 0.16 | 6.5 |
| *L. saxatilis* | Shell length | Aperture length | 0.73 | -0.14 | 7.3 |
| *L. saxatilis* | Shell length | Aperture thickness | 0.040 |  |  |
| *L. saxatilis* | Aperture length | Aperture thickness | 0.052 | 0.021 | 5.6 |

### 3.2 Predation experiments

In both the laboratory and field predation experiment, both crab species consumed all three species of snails. In the laboratory experiment, snail survival had a significant interactive response with crab species and snail species (GLM, *F*_2, 3534_ = 31.6, *p* < 0.01) and snail species and snail shell length (GLM, *F*_2, 3534_ = 21.1, *p* < 0.01). All snails smaller than 10 mm in shell length were consumable by both species but *H. sanguineus* could not consume 11 mm *L. obtusata* (Fig. 5). No snails died in the control treatment. In the field experiment, there was a significant interactive response to crab treatment, snail species, and snail shell length (GLM, *F*_4, 477_ = 2.5, *p* = 0.04). All crabs were capable of consuming all sizes of *L. saxatilis*, but *L. obtusata* had a size-refuge from *H. sanguineus* and *L. littorea* had a size-refuge from both crab species (Fig. 6). In control treatments, only three of 220 snails died (1.3%, one *L. obtusata* and two *L. littorea*). *Littorina saxatilis* had the highest proportion of its population at risk of consumption by all species of crabs (Fig. 6, > 55%), whereas *L. littorea* only had up to 11% of its population at risk. *Littorina obtusata* was intermediate with 33-48% of their population at risk of consumption.

**Figure 5.**
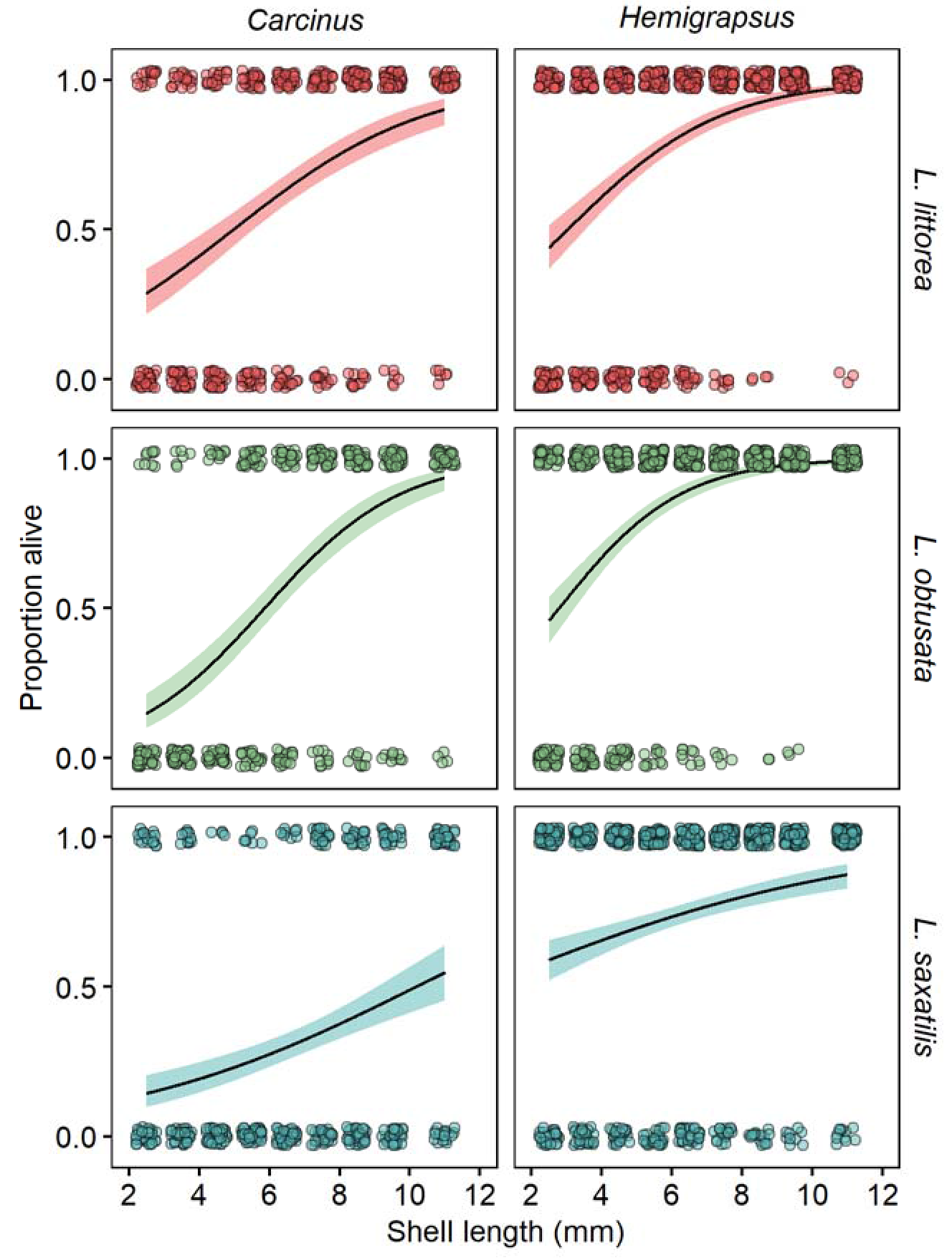
The effect of crab treatment and snail shell length on survival of three littorinid snails in laboratory enclosure experiments. Points are individual responses to the crab treatment. The solid line indicates the binomial regression fit and the colored areas are the 95% confidence intervals of those models.

**Figure 6.**
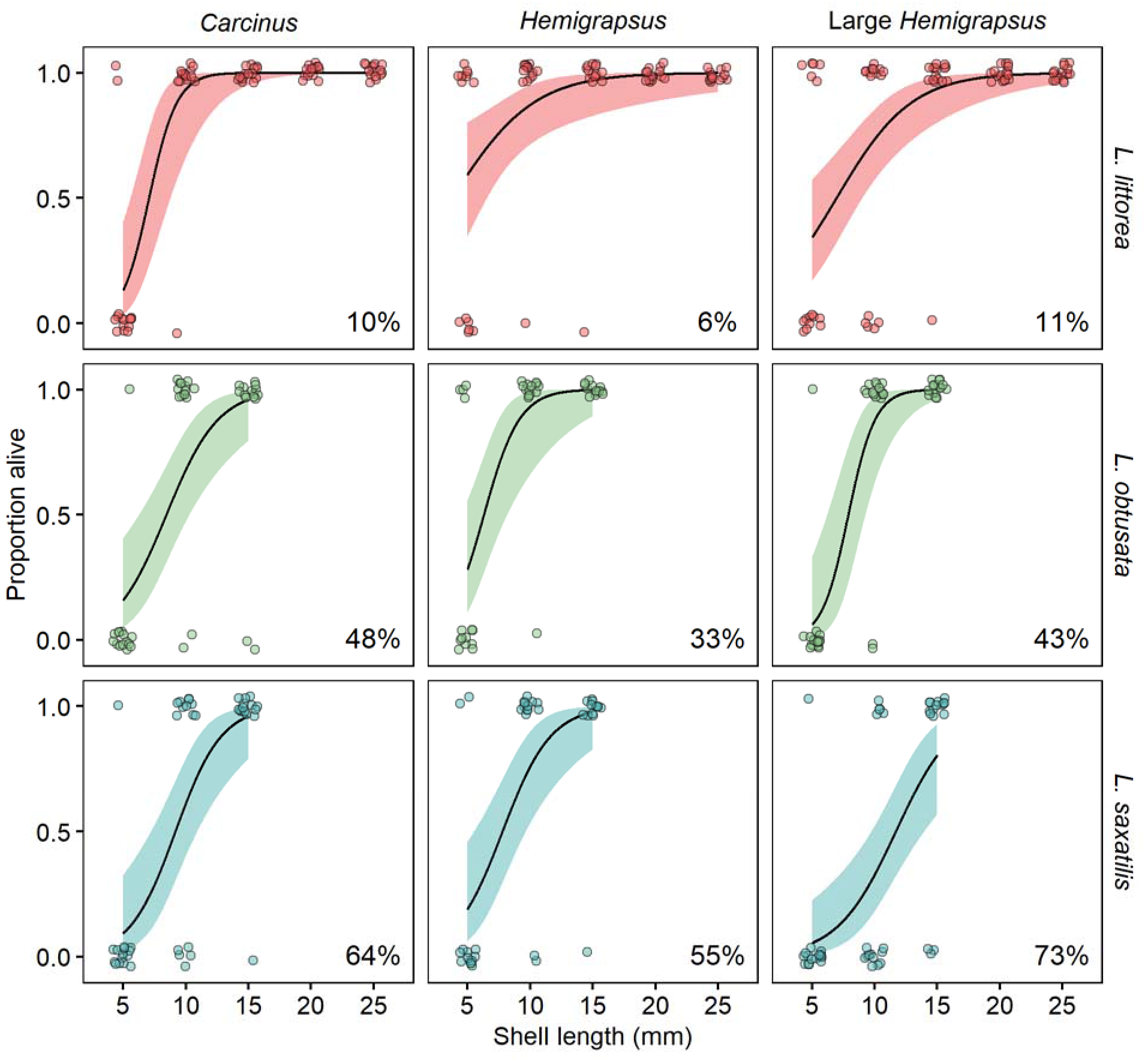
The effect of crab treatment and snail shell length on survival of three littorinid snails in field enclosure experiments. Points are individual responses to the crab treatment. The solid line indicates the binomial regression fit and the colored areas are the 95% confidence intervals of those models. Percentage of the population at risk of predation is reported in the bottom right of each panel.

## 4. Discussion

Distribution and morphology of native organisms can be altered by introduced species and this has been observed in littorinid snails with the historic invasions of *Littorina littorea* and *Carcinus maenas* (Behrens Yamada and Mansour, 1987; Trussell, 2000). Our study documents the extent of spatial and temporal overlap between littorinids and the two introduced intertidal crabs, *Hemigrapsus sanguineus* and *C. maenas*, both capable of exerting top-down pressure on snail abundances at all intertidal heights. Overall, the introduced crabs and *L. littorea* used a greater range of tidal heights than the native species, indicating that the introduced species use a broader range of resources than their native counterparts. Successful introduced species are more likely to be generalists than specialists, as species with broad tolerances are more likely to establish and spread in a novel environment and outcompete native species (Blackburn and Duncan, 2001).

Due to competitive pressure from *L. littorea, L. saxatilis* experienced reduced growth rates (Eastwood et al., 2006), and has been relegated to the high intertidal zone (Behrens Yamada and Mansour, 1987), which was once relatively free of crab predators. In regions where there are no crab predators and *L. Littorea* is absent, *L. saxatilis* can occupy the middle (Johannesson and Johannesson, 1990) to subtidal zones (Reid, 1996), and grows significantly larger in the lower intertidal (Behrens Yamada and Mansour, 1987). Our data suggests that prior to the introduction of *H. sanguineus*, *L.* saxatilis only came into contact with juvenile *C. maenas* in the high intertidal, but is now exposed to predation from adult *H. sanguineus* as well.

Our findings suggest that *L. saxatilis* may be under threat of extirpation in locations where *H. sanguineus* is becoming abundant. More than half of *L. saxatilis* are consumable by both introduced crabs and, unlike *C. maenas*, *H. sanguineus* has substantial overlap with *L. saxatilis* in the high intertidal. Additionally, *L. saxatilis*, unlike its congeners, increases in shell thickness linearly as it grows, and its shell remains thinner than the other species. All three snail species have similar initial investment in shell thickness (0.04-0.05 mm shell thickness per 1 mm shell length), but *L. littorea* and *Littorina obtusata* both more than double their shell thickness investment after reaching a critical point (20.9 and 7.5 mm, respectively). This shift may have been selected for in response to the increased predation pressure both species have experienced in the middle to shallow subtidal environments where there are significantly more decapod predators. Sustained predation pressure by crabs on a snail population often leads to shifts in shell morphology (reviewed in Reid, 1996). Snails living in areas with high predation pressure have thicker shells than those living in predator free areas (Chapman, 1997; Rochette et al., 2007; Teck, 2006). Additionally, sustained predation pressure on a population can cause a shift in shell morphology (Reid, 1996) as has been documented in *L. obtusata.* When *C. maenas* was introduced to the Gulf of Maine, flat-spired *L. obtusata* were selected for and contemporary populations now lack tall spires evident in older shell collections (Trussell, 2000).

Snails were most susceptible to predation at small sizes across all three snail species. Beyond 10 mm in shell length, predation susceptibility dropped substantially. For *L. obtusata* and *L. saxatilis*, a large proportion of the population was below 10 mm in length (67% and 86%, respectively) and therefore the population is at risk of predation from both introduced crabs for most of their life. *Littorina littorea* is larger bodied than the other two species and experiences a size-refuge for a substantial part of its life (Fig. 2). In addition, *L. littorea* also grows faster than *L. saxatilis* on a similar quality diet (Behrens Yamada and Mansour, 1987), thus moving more quickly out of the vulnerable stage. While *L. saxatilis* is more vulnerable to crab predation than the other snails, it may be the best able to locally adapt to novel pressures. *Littorina saxatilis* has direct-developing larvae (Johannesson, 2003) and has high location-fidelity as an adult, moving only a few meters over the course of several months (Janson, 1983). Combined, these factors lead to distinct localized ecotypes (Johannesson, 2003; Reid, 1996), which allow morphology to differ drastically across small distances (Johannesson and Johannesson, 1990).

Here, we evaluate the effects of the ongoing invasion of *H. sanguineus* on littorinids in the Gulf of Maine, particularly its effect on *L. saxatilis. L. littorea and L. obtusata* are able to withstand predation pressure from crab species through thickening their shells and growing quickly to reach a size refuge. However, *L. saxatilis* has a small body size, a relatively thin shell, and spatially overlaps with the new crab predator *H. sanguineus*. It has already been forced into the high intertidal by competition with *L. littorea* (Behrens Yamada and Mansour, 1987), where there are lower quality food sources, and environmental conditions are more extreme. There is also the potential for additional predators or competitors, added through range expansions or new invasions, which are taking place at an accelerated rate in the Gulf of Maine. Abiotic conditions are becoming more extreme with the progression of climate change, which may make the high intertidal a much harsher environment. While *L. saxatilis* possesses the ability to locally adapt, it is unclear whether it will be able to keep pace with its rapidly changing environment or if competition, predation, and climate change will push populations of *L. saxatilis* to local extirpation.

## Acknowledgements

We would like to thank the University of New Hampshire for use of their laboratory space. Larry Harris provided guidance, laboratory space, and supplies.

## Funding

This research did not receive any specific grant from funding agencies in the public, commercial, or not-for-profit sectors.

## Notes

### Competing Interest Statement

The authors have declared no competing interest.

